# The effect of microbial selection on the occurrence-abundance patterns of microbiomes

**DOI:** 10.1101/2021.09.02.458693

**Authors:** Román Zapién-Campos, Michael Sieber, Arne Traulsen

## Abstract

Theoretical models are useful to investigate the drivers of community dynamics. Notable are models that consider the events of death, birth, and immigration of individuals assuming they only depend on their abundance – thus, all types share the same parameters. The community level expectations arising from these simple models and their agreement to empirical data have been discussed extensively, often suggesting that in nature, rates might indeed be neutral or their differences not important. But, how robust are these model predictions to type-specific rates? And, what are the consequences at the level of types? Here, we address these questions moving from simple to diverse communities. For this, we build a model where types are differently adapted to the environment. We adapt a computational method from the literature to compute equilibrium distributions of the abundance. Then, we look into the occurrence-abundance pattern often reported in microbial communities. We observe that large immigration and biodiversity – common in microbial systems – lead to such patterns, regardless of whether the rates are neutral or non-neutral. We conclude by discussing the implications to interpret and test empirical data.

## 1 Introduction

Theoretical models have been instrumental in understanding ecological systems. Historically, a handful of puzzling natural observations have motivated their development – from the limits of exponential growth by Malthus [1] to the competition of species by Lotka and Volterra [2, 3].

The stark difference of the frequencies of species within communities is one such observation. While few species are very abundant, many others barely appear in community surveys [4]. Two hypotheses have dominated the scientific discussions. On one hand, it is proposed that biotic interactions and environmental filtering make trophically similar species occupy different niches, which allows differences in abundance while preserving diversity. This is known as niche theory [5]. Alternatively, Hubbell and others [6] have emphasized that even if niche differences are discounted, so only species’ abundances matter, random fluctuations can lead to the patterns of abundance and diversity observed in nature. This is known as neutral ecological theory [7].

Despite their stringent assumptions, neutral models often predict patterns observed in communities as different as the tropical rainforest of Barro Colorado island [7] and host-associated microbiomes [8, 9, 10]. With time, neutral models have become null hypotheses used to discard the need for complex mechanistic explanations in data at the community level [6].

But how does a neutral model work? In a neutral model the death and birth of individuals account for changes in community composition. However, because each rate is identical for all types, after some time, stochastic drift leads to the extinction of all but one type [11]. Thus, to preserve diversity, an external source of individuals by immigration or speciation is needed. Here, neutral theory builds upon island biogeography. In this theory, MacArthur and Wilson [12] have modelled the community composition of small habitats (“islands”) connected by migration to a larger habitat (“mainland”). In neutral models, a local community commonly receives individuals from an external and larger community [13]. Such community can itself undergo internal changes or, by separation of time scales, assumed to be constant [7, 13].

Early on, neutral models have been used in macroecology to address the patterns of diversity and abundance of species [7, 6]. More recently, driven by developments in sequencing technologies, the study of patterns of occurrence and mean frequency in microbial communities has become possible [14]. At this scale, ecological drift also seems to greatly influence the community dynamics, leading to hypothesize that many microbial taxa could be classified as neutral [15, 10]. However, few taxa, referred to as non-neutral, have occurrences and frequencies different than neutrally expected. It has been suggested that the last group might include, among others, pathogens and symbionts [10].

At least two possibilities could lead to deviations from neutrality. Either different processes from those in the neutral model are necessary, or, alternatively, not all the parameters of the model are actually neutral. Both of these lead to develop models of selection [11]. Although many such models have been developed from niche theory assumptions, fewer have been developed from a neutral theory basis [6, 16]. A direct connection from neutral to selective models would allow to comparing their patterns while acknowledging that both might be operating simultaneously. Indeed, the role of non-neutral processes can only be rejected after ensuring that they can not produce “neutral” patterns [16], especially in data.

Neutral and niche models have been connected in several ways [17, 18, 13]. Some authors have assumed that the rate of types are solely determined by the environment, finding that neutrality might overshadow the niche structure effect [19], depending on diversity, dispersal, and niche overlap [17]. Alternatively, using Lotka-Volterra models with immigration, the effect of competitive interactions has been studied. Early models focused on intraspecific [20] or interspecific [21] competition. Later on, both were considered simultaneously. Haegeman and Loreau tuned the niche overlap using symmetric interactions to investigate the success behind the neutral assumption [18]. Kessler and Shnerb classified the dynamics emerging from interspecific interactions, finding that the neutral case links all classes [13]. Focusing on intraspecific interactions, Gravel et al. studied the influence of immigration, suggesting a continuum from competitive to stochastic exclusion [17]. Throughout these studies, diversity, community size, and environmental fluctuations seem to have great relevance, as pointed out by Chisholm and Pacala and Fisher and Mehta.

This previous research has proven useful to bridge neutral and selective theories. The link has been instrumental to consider migration, speciation, and stochastic demography key components in ecology. Along this line and motivated by the particularities of microbial communities, large community size and taxa diversity [15, 23, 14], here we investigate the commonly observed occurrence-abundance pattern in neutral and non-neutral contexts. Similarly to Sloan et al. and Allouche and Kadmon, we model death, birth, and immigration within a community, but in contrast to these neutral models, type-specific growth and death rates are determined by the environment.

## 2 Results

### 2.1 A spatially-implicit death-birth model with immigration

We consider a set of local communities connected by immigration to a larger community which contains multiple types of individuals. While local communities change as a result of the death, birth, and immigration of individuals, the larger community changes on a much longer time-scale – so immigration to local communities can be assumed to be constant. To derive a dynamical equation of a local community composition, we account for the events that change the frequency *x_i_* of each type *i* = 1, …, *S* within each local community. Individuals die with a rate proportional to the product *x_i_ϕ_i_* of their frequency and their death rate *ϕ_i_*. Additionally, they are born proportional to the product *x_i_f_i_* of their frequency and their growth rate *f_i_* – or arrive with a fraction of the immigration rate *m* that reflects their frequency *p_i_* in the external environment. Combining these processes, we obtain

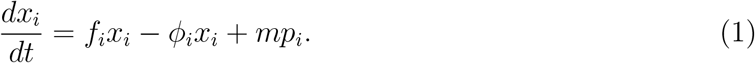

Assume for now an equal death rate for all types, *ϕ_i_* = *ϕ*, so only *f_i_*, *m*, and *p_i_* are free parameters. To hold the community size constant, we use ∑_*i*_ *dx_i_*/*dt* = 0 to find 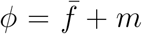, where 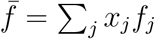 is the average growth rate of a randomly selected individual. In this way

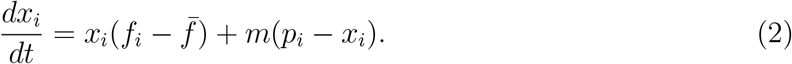

Without immigration, *m* = 0, Eq. (2) shows that only types whose growth rate is larger than the average increase. After sufficient time, only the type with the largest growth rate remains. Coexistence is only possible in the neutral case, where all types have the same growth rate, 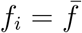. There, the initial frequencies remain unchanged. Immigration, *m* > 0, creates an equilibrium that resembles the external composition, *p_i_*, that for sufficiently large immigration might promote coexistence, especially if types with small growth rate migrate more. Similar results are obtained if we assume equal growth rate for all types, *f_i_* = *f* in Eq. (1) instead. In the general case, growth and death rates have opposing effects.

Eq. (1) provides useful insights about the dynamics and equilibria; however, only a stochastic model would allow us to compute observables such as the occurrence frequency and the variance. To develop such a model, we track the vector of absolute abundances instead, **n**, and list the transition rates that change it. The increase of type *i* by one individual occurs at the expense of the decrease of type *j*,

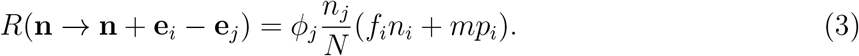

Here **e**_*i*_ and **e**_*j*_ are vectors whose i-th or j-th element equals one and zero elsewhere. The carrying capacity of the community is given by *N*. The master equation accounts for changes in the probability of observing the community composition **n** through time,

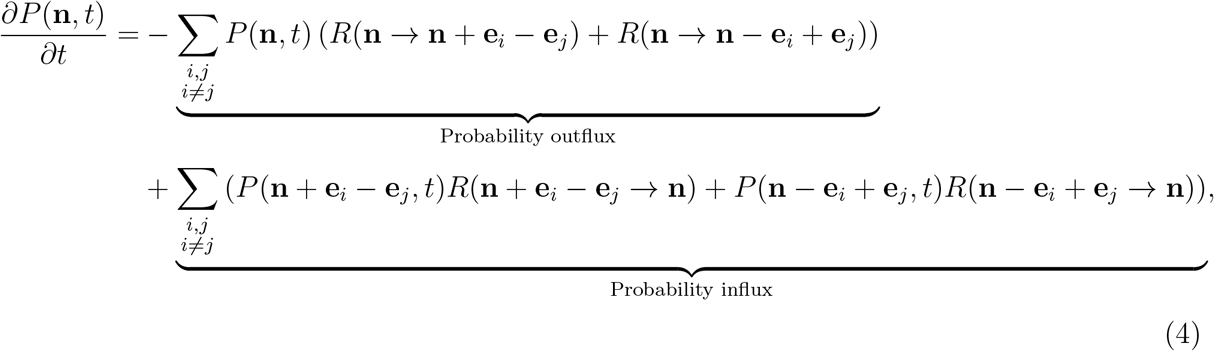

where *P*(**n**, *t*) is the probability density of community composition **n** at time *t*.

In this work, we investigate the probability distribution at equilibrium, i.e. the state where the master equation equals zero. In this case, the influx and outflux to each state balance each other, ending up with a system of equations that can be solved to find *P*(**n**). For communities composed of two types (*S* = 2), a detailed balance analysis [24] leads to a recurrence equation of the arbitrarily denoted type 1

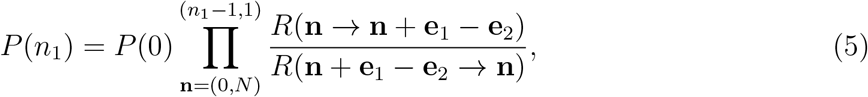

satisfying 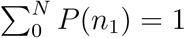. In this case, the transition rates are simplified to the single variable *n*_1_, using *n*_2_ = *N* – *n*_1_ and *p_2_* = 1 – *p*_1_.

For communities with more than two types (*S* > 2) analyses are more challenging, as all possible compositions must be considered. This is particularly true for microbial communities, where many types interact (10^1^ to 10^4^ taxa are common) in large communities (10^3^ to 10^14^ individuals). Although a recurrence equation exists [21], the exponential increase in the number of states and transitions with *S* and *N*, make its computation unfeasible. This is a problem common to microscopic and even mesoscopic descriptions, which has been deemed “the curse of dimensionality” [25]. In neutral models, the equality of rates allows to reduce analyses to a single dimension – that of a focal type [15]. However, unless density dependence is neglected, non-neutral models are inherently multidimensional, as transitions depend on the current community composition.

A potential way forward is to acknowledge that, typically, rather than being interested in the probability of every possible community, we are interested in marginal probabilities. In other words, the added probabilities over various dimensions. Methods of model reduction have been developed towards this aim. Based on various assumptions, these methods sacrifice “microscopic” information in the interest of specific observables. Jahnke introduced the *model reduction by conditional expectations* (MRCE), where, while selected types are described stochastically, others are modeled using a mean-field approximation [26]. The MRCE is derived from the Bayes theorem, by which *P*(**n**, *t*) is given by the product of two probabilities, one for some chosen types and one for the conditional probability of the others. Then, the probabilities of the others are replaced by expected abundances. Because of the last point, the method is particularly suited to systems where types have peaked distributions and large populations – a situation that can be akin to some microbial communities.

In this paper we combine the MRCE method [26] with a detailed balance analysis [24] to compute the marginal probability distribution of types within a microbial community. For each distribution at equilibrium, we extract the probability of occurrence, *P*(*n_i_* ≥ 1), the mean frequency *E*(*n_i_*)/*N*, and compare them in situations of neutrality versus non-neutrality.

To apply the MRCE method, we adapt our model to the convention in [26]. First, we split the vector of abundances 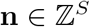 into a focal type *i*, *n_i_*, and the set of others, 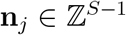, *j* ≠ *i*, for which the marginal probability, 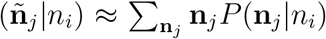, and the expected abundance conditioned on the focal type, (**ñ**_*j*_|*n_i_*) ≈ ∑_**n**_*j*__ **n**_*j*_ *P*(**n**_*j*_|*n_i_*), are approximated. Then, each transition rate is factored as the product of rates of the focal type and other types,

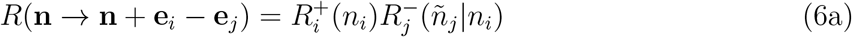

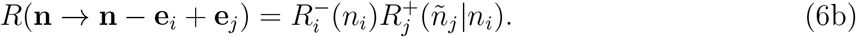

In our model, 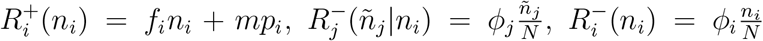 and 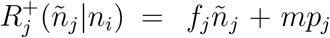. With these transformations, the equilibrium is given by the simplified master equation of the focal type *i*,

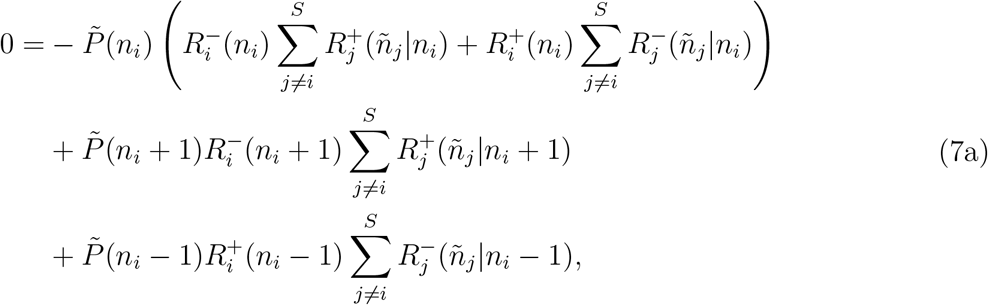

and a set of equations for the expected abundance of the others conditioned on the abundance of the focal type (**ñ**_*j*_|*n_i_*),

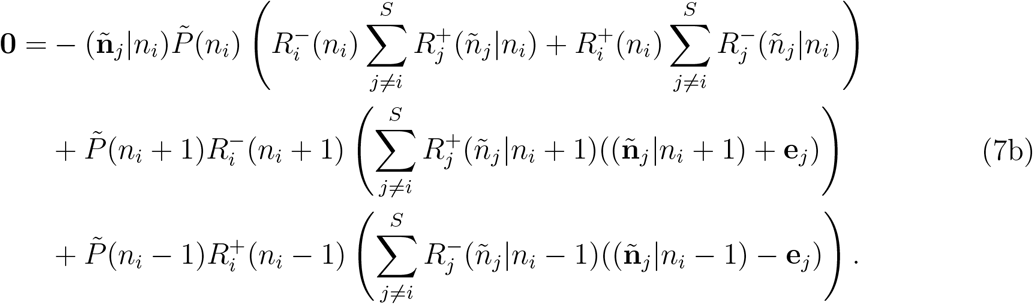

We solve this system of equations in the range of *n_i_* = 0, …, *N*, starting from *n_i_* = *N*. By definition 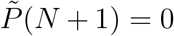, so no probability flux to or from *N* + 1 occurs. Then, the influx from *n_i_* = *N* implies 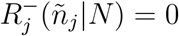, specifically (**ñ**_*j*_|*N*) = **0**. We end up with a simplified system of equations for *n_i_* = *N*. To compute 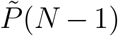 and (**ñ**_*j*_|*N* – 1) from this, we assume without loss of generality 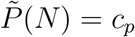, where *c_p_* is a positive constant. Consecutive 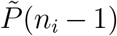 and (**ñ**_*j*_|*n_i_* – 1) are computed iteratively. Finally, the normalization 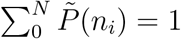 is enforced.

A reliable numerical method is needed to solve Eq. (7a-7b). The large difference between the magnitudes of 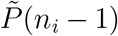 and (**ñ**_*j*_|*n_i_* – 1) can cause numerical problems. To avoid them, we extract 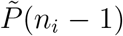 from Eq. (7a) and substitute it in Eq. (7b) – note that all else are known values. The resulting system of equations is solved for (**ñ**_*j*_|*n_i_* – 1), and these substituted in Eq. (7a) to compute 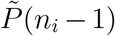. Caution is needed in cases that lead to a normalized 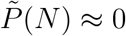, especially if computations are performed in a machine with limited float representation. In this case, we find the 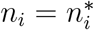 closest to *n_i_* = *N* that while declaring 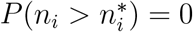 and 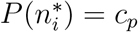 allows for the iterative solution.

Compared to the fully stochastic model that scales with 2^−*S*^*N^S^*, here, we solve *N*(*S* – 1) equations for the marginal probability of each type, i.e. *N*(*S*^2^ – *S*) equations for the community. This model reduction allows us to approximate the equilibrium of large communities with many interacting types more rapidly.

### 2.2 The neutral expectation

We start by considering the neutral case – a situation where the rates of all types are equal (*f_i_* = *ϕ_i_* = 1 for all *i* in {1, …, *S*}). In contrast to the deterministic model at equilibrium, Eq. (2), the frequencies of single stochastic realizations change through time, driven by the probabilistic nature of events. As a result, a distribution of frequencies centered at the value set by the source of immigrants (*p_i_*) emerges. The spread of this distribution inversely depends on the magnitude of the immigration, *m*.

As shown in Fig. 1A, large immigration drives the equilibrium distribution towards its mean value, *p_i_*. On the contrary, without or little immigration, the distribution splits. Thus, the frequencies zero (no individuals of the i-th type) and one (only individuals of the i-th type) are the most probable, decaying towards intermediate frequencies. This is a consequence of noisy fluctuations that, for a single realization, lead to the extinction of all but one type. Whether the frequency one or zero is most likely depends on the proximity of the initial state.

**Figure 1:**
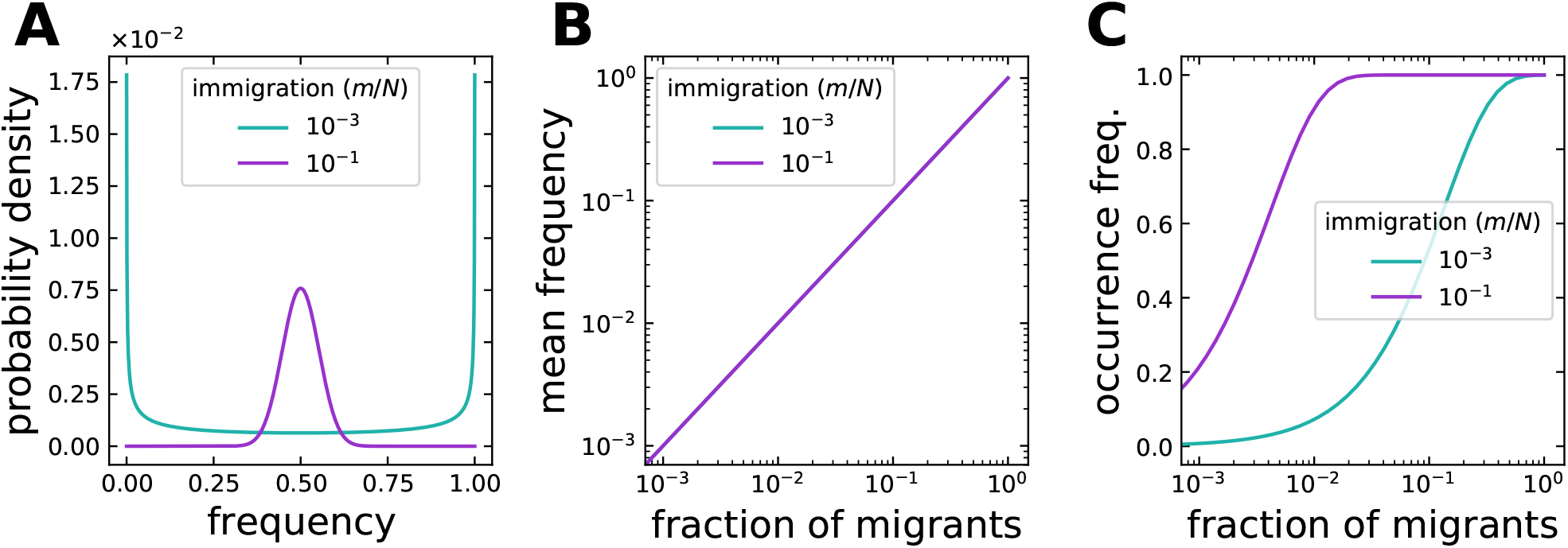
Expected equilibrium of a type if rates in the community are neutral. (**A**) If the immigration is very small, the population either goes extinct or reaches fixation. A larger immigration reduces the variation in frequency, centered at its fraction of immigrants (here *p*_1_ = 0.5). (**B**) The mean frequency increases with the fraction of immigrants *p*_1_, but is independent of the immigration rate *m*. (**C**) Also the occurrence frequency increase with the fraction of immigrants (*p*_1_), but in an S-shaped manner that depends on *m*. Deviations from these patterns have been suggested to indicate non-neutral rates [10]. The community size is *N* = 10^3^.

The mean frequency of the stochastic model identically corresponds to the frequency of the deterministic model. As shown in Fig. 1B, regardless of the total immigration, the mean frequency of a type increases linearly with the fraction of migrants of its kind.

Besides the mean frequency, one of the simplest, but most informative observables is the occurrence frequency of individuals of a given type in the community. In other words, the probability of observing at least one individual of that type, *P*(*n_i_* ≥ 1). Immigration increases this probability up to the point where the type is always observed in the community (Fig. 1C). Importantly, this probability does not increase linearly with the fraction of migrants. Instead, an S-shaped curve is observed, where changes of immigration of rare or abundant types do not modify their occurrence.

Using two simple observables, the mean frequency and the occurrence frequency, we can describe the state of types within a community. In the following, we relax the assumption of neutrality – not enforcing equal growth and death rates. Then, we contrast both observables to their neutral expectation.

### 2.3 Immigration lessens the effect of growth and death differences

To understand the effect of non-neutral rates, we start from a community composed of only two types. Furthermore, we assume only one of them has a non-neutral rate, either *f_i_* or *ϕ_i_*. In this way, we aim to see the effect in the neutral and non-neutral fractions of the community.

For a growth rate below one (*f*_1_ < 1) or a death rate above one (*ϕ*_1_ > 1), the non-neutral type has a reduced mean frequency that preserves its linear relationship to the fraction of immigrants (Fig. 2A-B and Fig. 3A-B). However, in contrast to the neutral expectation, immigration does play a role, as large migration can reduce the changes occurring in the internal community dynamics (compare panels A to B in Fig. 2-3). In this context, the neutral type (*f*_2_ = *ϕ*_2_ = 1) benefits from the reduced proliferation of its partner, thus, gaining in frequency, especially if most immigrants belong to the neutral type.

**Figure 2:**
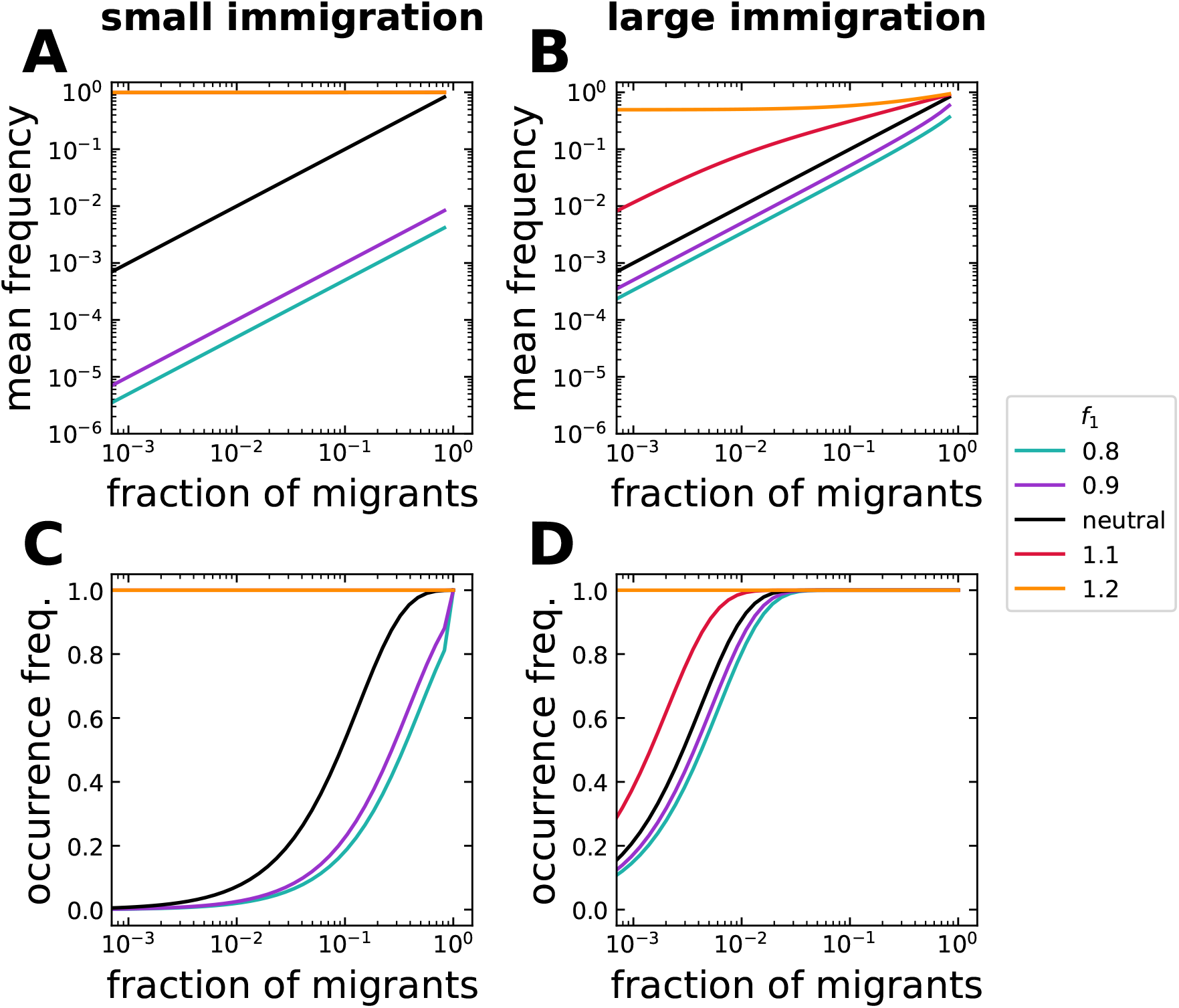
Effect of non-neutral growth rates on the equilibrium of a community with two types. One of two types has non-neutral growth rate (*f*_1_ ≠ *f*_2_ = 1), but the death rate is neutral (*ϕ*_1_ = *ϕ*_2_ = 1). In contrast to its all-neutral (*f*_1_ = *f*_2_ = 1) expectation, a lower growth rate of the non-neutral type (*f*_1_ < *f*_2_) reduces its mean frequency and occurrence. The change can be of several orders of magnitude. Inversely, a larger growth rate of the non-neutral type (*f*_1_ > *f*_2_) increases its mean frequency and occurrence. The effect of growth rate differences on the internal dynamics is reduced if immigration is larger, especially for slowly growing types. Immigration is (**A**, **C**) *m/N* = 10^−3^ and (**B**, **D**) *m/N* = 10^−1^, with community size *N* = 10^3^.

**Figure 3:**
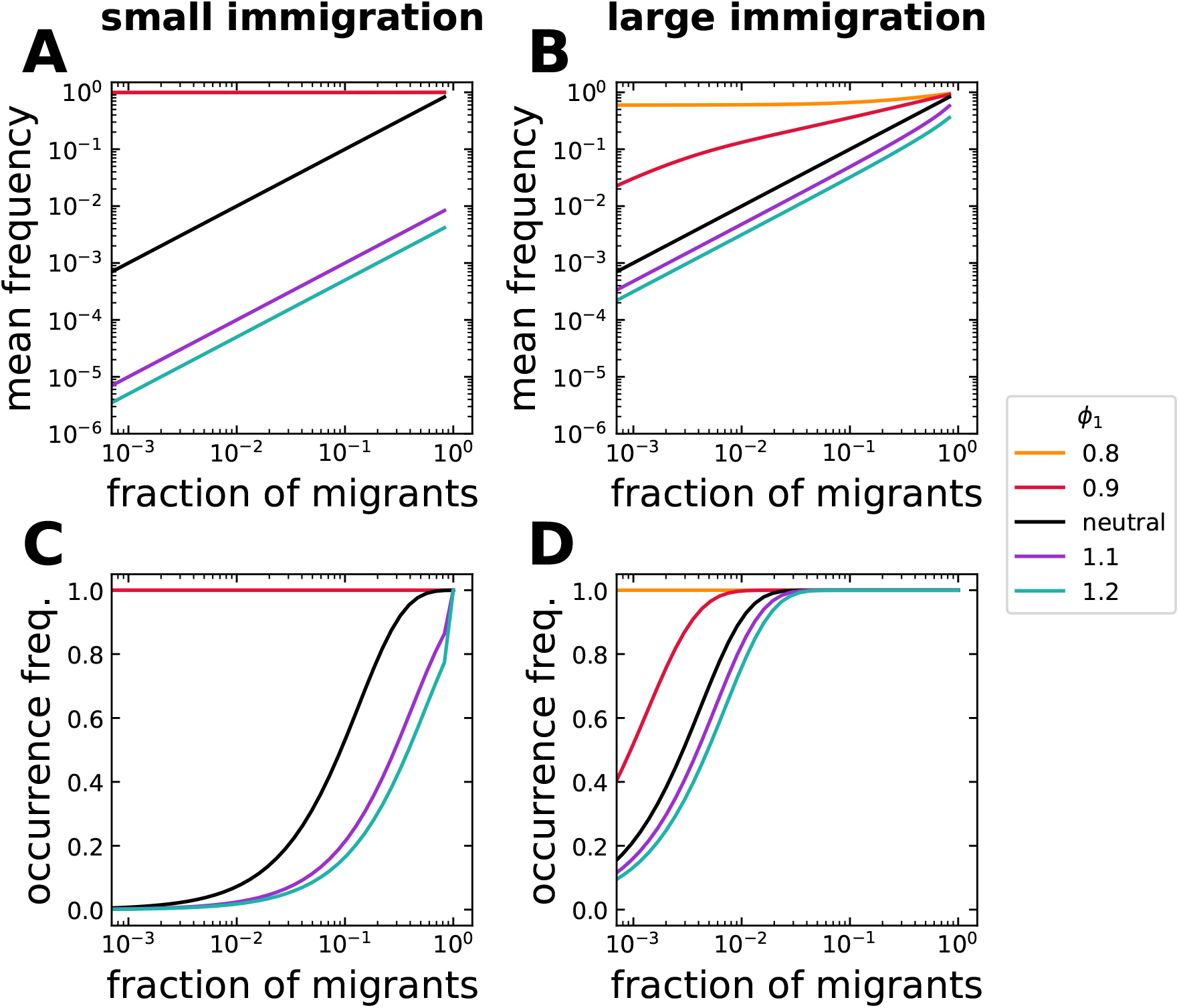
Effect of non-neutral death rates on the equilibrium of a community with two types. One of two types has non-neutral death rate (*ϕ*_1_ ≠ *ϕ*_2_ = 1), but neutral growth rate (*f*_1_ = *f*_2_ = 1). Differences in death rates modify the mean frequency and occurrence of both types. A larger immigration reduces differences to the all-neutral (*ϕ*_1_ = *ϕ*_2_ = 1) expectation in a similar fashion to differences in growth rate (Fig. 2). Immigration is (**A**, **C**) *m/N* = 10^−3^ and (**B**, **D**) *m/N* = 10^−1^, with community size *N* = 10^3^.

A similar picture arises for the occurrence pattern. While the non-neutral type occurs less frequently, the neutral type thrives, occurring more often than when both types are neutral (Fig. 2C-D and Fig. 3C-D). The change can be as severe as losing all non-neutral individuals from the community (panel C in Fig. 2-3). Crucially, large total immigration can prevent this (compare panels C to D in Fig. 2-3), even if most migrants are of the neutral type.

Once the roles are reversed, so the non-neutral growth rate is above one (*f*_1_ > 1) or the death rate below one (*ϕ*_1_ < 1), the mean frequency and occurrence patterns mirror the previous results (Fig. 2 and Fig. 3). Although changes produced by non-neutrality in growth (*f*_1_ ≠ 1) or death (*ϕ*_1_ ≠ 1) rates are qualitatively similar, they show quantitative differences.

We conclude that even for the simplest community (one with two types), just one non-neutral rate is enough to change the community occurrences and abundances substantially from their all-neutral expectation. This is more visible through the mean frequency (as changes of several orders of magnitude are possible) and for communities with little external migration – where the internal dynamics is more important.

### 2.4 Neutral and non-neutral patterns are similar at the community level but full of differences at the level of types

Communities with two types might occur *in vitro*. However, in nature, communities are much more diverse, especially for microbes. We have produced random instances of such diverse communities, sampling growth and death rates, *f_i_* and *ϕ_i_*, from a normal distribution with mean one and a desired standard deviation. Similarly, we have produced random fractions of migrants, *p_i_*, just conditioned on the Gini index of the community,

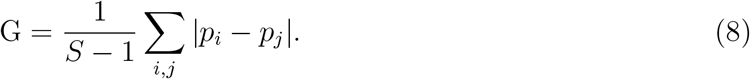

This number that indicates the asymmetry in immigration between types from zero to one, allow us to compare communities quantitatively, regardless of their number of types *S*. As an example, for *G* = 0 the fractions of migrants are identical for each type, while for *G* = 1 the source pool only contains a single type.

Using these parameters, we have computed the occurrence and abundance frequency of all types in a certain community. Interestingly, the community patterns that we observe are very similar to those expected from neutrality (Fig. 4A compared to Fig. 1C) – even if asymmetries of growth, death, and immigration increase (Fig. 5). In particular, large immigration together with high biodiversity consistently lead to these patterns (Fig. 5). This indicates that neither neutrality nor non-neutrality, but large immigration and biodiversity are behind these patterns.

**Figure 4:**
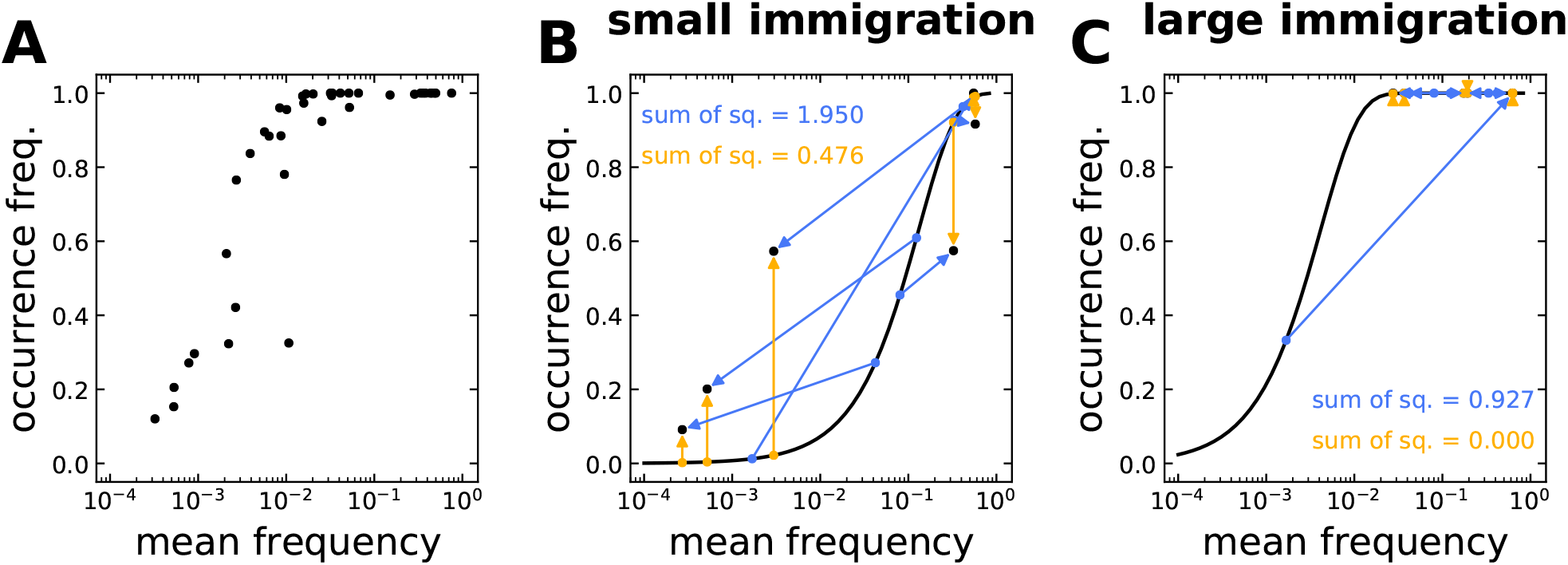
Occurrence-abundance pattern in general non-neutral communities. (**A**) The non-neutral pattern of a diverse community largely resembles neutral patterns, see Fig. 1C. (**B**) However, the change from neutrality of each type can be large (blue arrows), shown here for *m/N* = 10^−3^. In general, the mean frequency does not equal the fraction of immigrants *p_i_*, assuming otherwise underestimates the change from neutrality (yellow arrows). (**C**) Similar to a community with two types, Fig. 2-3, the overlap to the neutral expectation increases when immigration, *m*, is increased to *m/N* = 10^−1^. The growth and death rates, *f_i_* and *ϕ_i_*, were sampled from a normal distribution with mean 1 and standard deviation 0.1, where *P*(*f_i_* < 0.8) = *P*(*f_i_* > 1.2) ≈ 0.023 and *P*(*ϕ_i_* < 0.8) = *P*(*ϕ_i_* > 1.2) ≈ 0.023. The fractions of migrants *p_i_* range from 10^−4^ to 10^−1^ and have a *G* ≈ 0.6, Eq. (8), indicating intermediate immigration asymmetry. Except from the immigration rate *m*, all rates in (**B**-**C**) are equal. The community size is *N* = 10^3^.

**Figure 5:**
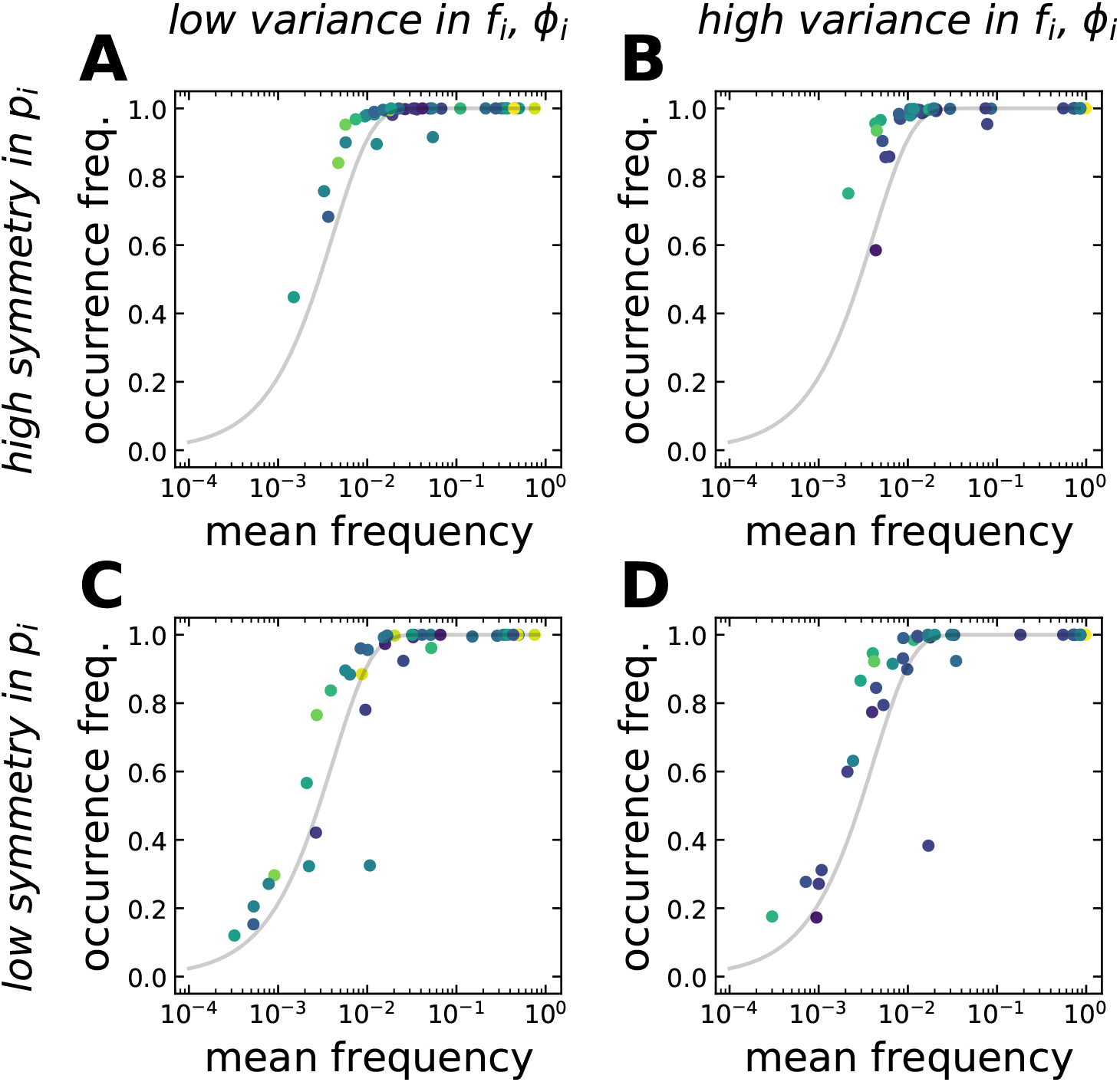
Occurrence-abundance pattern for different levels of asymmetry in the parameters. **A** pattern is robust to various asymmetries in the immigration, *p_i_*, and growth and death rates, *f_i_* and *ϕ_i_*. Each community has forty types. For low symmetry in immigration, the types span the range more widely. Colors from dark to light indicate how non-neutral a type is, quantified as the geometric distance from (*f_i_*, *ϕ_i_*) = (1, 1). Types overlap regardless of their non-neutrality. The fractions of immigrants, *p_i_*, have a *G* ≈ 0.3 (**A**-**B**) or *G* ≈ 0.6 (**C**-**D**). The growth and death rates, *f_i_* and *ϕ_i_*, were sampled from a normal distribution with mean 1 and standard deviations 0.1 (**A**, **C**) or 0.2 (**B**, **D**). In the last case, *P*(*f_i_* < 0.8) = *P*(*f_i_* > 1.2) ≈ 0.159 and *P*(*ϕ_i_* < 0.8) = *P*(*ϕ_i_* > 1.2) ≈ 0.159. Immigration is *m/N* = 10^−1^, with community size *N* = 10^3^.

Even when neutral and non-neutral patterns are similar at the community level, we observe large differences at the level of types. While in the “all-neutral” case, the mean frequency equals the fraction of migrants, *E*(*n_i_*)/*N* = *p_i_*, this is not the case in a non-neutral scenario (Fig. 4B-C). Neither is for the occurrence frequency. The distance from the neutral expectation of each type is not simply related to the level of non-neutrality of its own parameters. Rather, neutral and non-neutral types fall on, above, or below the neutral expectation (Fig. 5), highlighting the inherent multidimensionality determining the equilibrium of these communities.

To investigate the effect of single parameters at the level of types, we chose two representative types – one close to the neutral expectation, and another one distant from it (Fig. 6A). Our results show that types do not remain on or far from the neutral expectation. Rather, the relative magnitude of their growth and death rate, *f_i_* and *ϕ_i_*, is crucial to observe simultaneous decrease or increase in occurrence and mean frequency (Fig. 6C-D). In particular, types with a smaller fraction of immigrants, *p_i_*, experience more abrupt changes. Only large fractions of immigrants allow to overcome the effect of growth and death rate differences, leading to large occurrence and mean frequency at the level of types (Fig. 6B).

**Figure 6:**
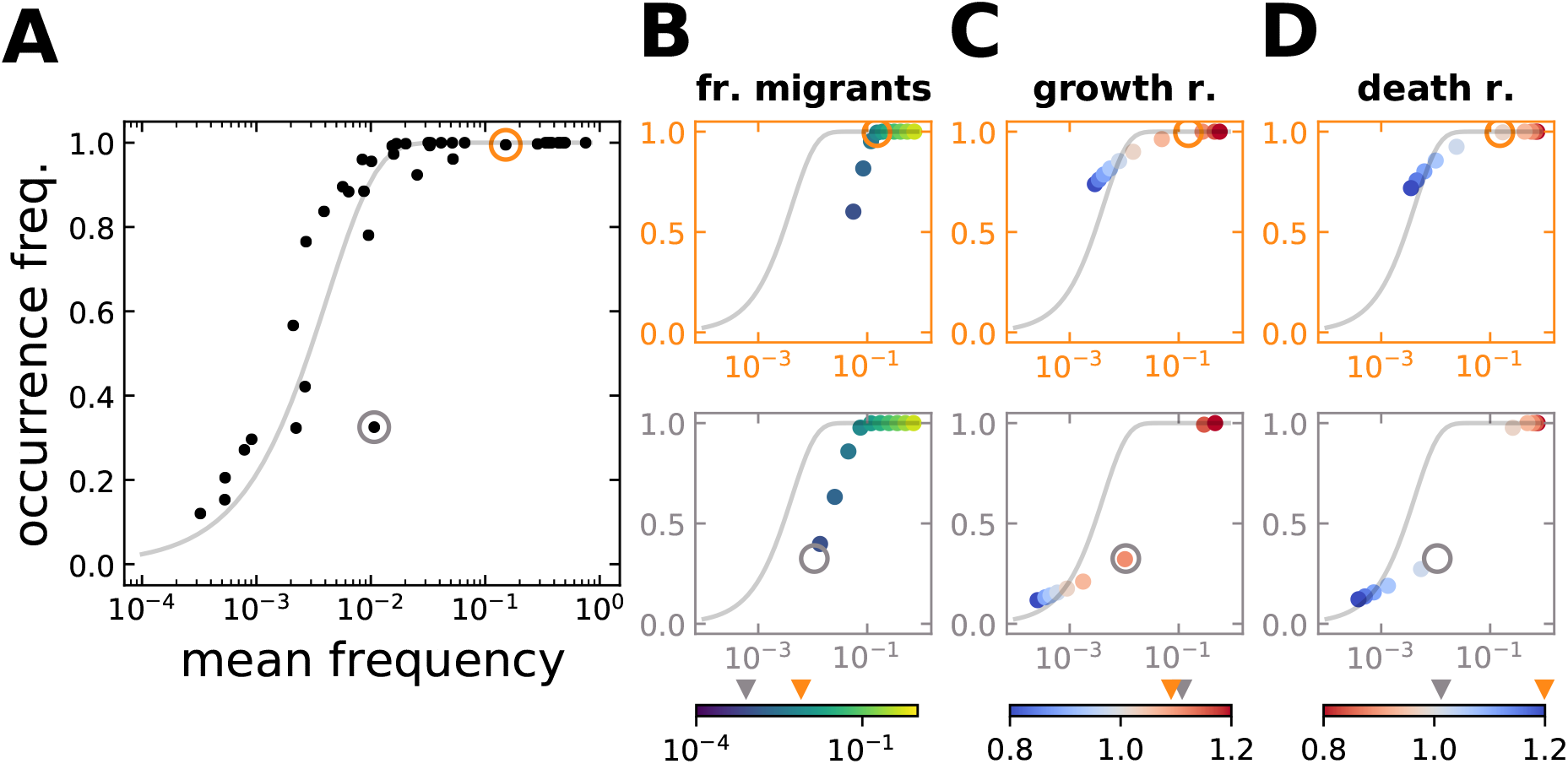
Effect of growth, death, and immigration at the level of types. (**A**) The community shown corresponds to Fig. 5C, with *G* ≈ 0.6 for *p_i_*, and *f_i_* and *ϕ_i_* drawn from 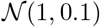. Two types are spotted by circles, one that falls on the neutral expectation and the other distant from it. Single parameters are modified in (**B**-**D**) for both types. Arrows in the colorbars indicate their original values. (**B**) For large fractions of migrants, *p_i_*, non-neutral types are indistinguishable from the neutral expectation; only for small fractions they are below it. (**C**) Different growth rates, *f_i_*, lead non-neutral types to fall on, above, or below the neutral expectation. Changes are especially abrupt for the type with less immigration. (**D**) Different death rates, *ϕ_i_*, mirror the effect of changing growth rates qualitatively. Immigration is *m/N* = 10^−1^, with community size *N* = 10^3^.

### 2.5 To test neutrality the niche structure must be known first

So far we have used our model to compute observables based on known parameters. However, we can invert this process to infer parameters from simulations or experimental data.

Particularly relevant is the possibility of testing niche structure in data [15, 8, 9, 10]. Our model indicates care is needed to quantify the true difference from neutrality (Fig. 4B-C). In fact, the comparison of the selective case to the neutral case can only be inferred after fitting all parameters of the general model (*m*, *p_i_*, *f_i_*, and *ϕ_i_* for all *i*). This is in contrast to the – often used – method by Sloan et al. for neutral conditions, where only the immigration rate *m* is fitted, while all growth and death rates are assumed *f_i_* = *ϕ_i_* = 1, and the fraction of migrants *p_i_* equalled to the mean frequency *E*(*n_i_*)/*N*. Our results indicate these assumptions on the data are unfounded and lead to underestimate niche structure (Fig. 4B-C), especially in large communities with many types. Moreover, the consistent occurrence-abundance pattern that we observe (Fig. 5), and often reported in data [8, 9, 10], emerges from a general death-birth processes with immigration, Eq. (3), not just from a neutral process (where *f_i_* = *ϕ_i_* = 1 for all *i*). Niche structure – and thus neutrality – can not be discarded or confirmed if certain parameters are fixed *a priori* [15].

The large number of parameters to be fitted requires large datasets. For a community with *S* types, 3*S* + 1 parameters must be fitted, thus requiring at least 3*S* + 1 data points. The 2*S* data points obtained from the occurrence and mean frequencies are not sufficient. We propose to include additional observables that can be readily computed from data [27]. These might include, but not be limited to, raw and central moments of the frequency. From this set of observables, available Bayesian methods [28] can be used to infer the parameters using Eq. (7a-7b).

In Fig. 7, we show two potential observables, the variance and the second moment of the distribution. In a community with two types, *S* = 2, both observables reflect the differences in growth, *f_i_*, and death rates, *ϕ_i_*. Only some variances overlap for distinct rates. In this sense, the second raw moment might provide more information to discriminate them. A set of similar observables could allow to characterize the rates of empirical communities.

**Figure 7:**
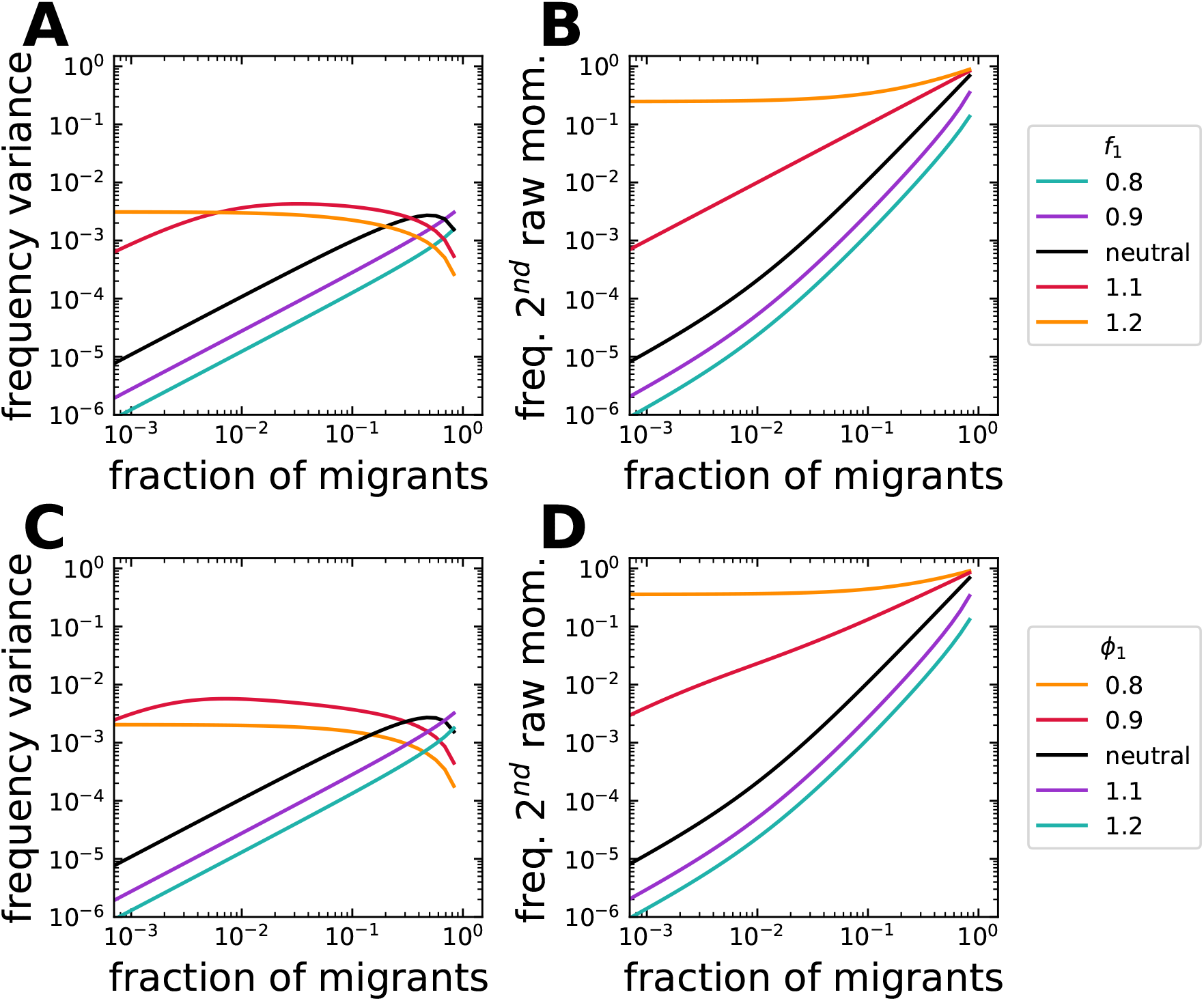
Variance and second raw moment of the frequency. A community with two types is considered. (**A**-**B**) One type has non-neutral growth rate (*f*_1_ ≠ *f*_2_ = 1) but neutral death rate (*ϕ*_1_ = *ϕ*_2_ = 1), or (**C**-**D**) a neutral growth rate (*f*_1_ = *f*_2_ = 1) but non-neutral death rate (*ϕ*_1_ ≠ *ϕ*_2_ = 1). (**B**, **D**) *f*_1_ > 1 and *f*_1_ < 1 lead to a second raw moment above or below the neutral expectation, respectively. This moment increases continuously with the fraction of migrants, *p_i_*, while the variance reaches a maximum at intermediate *p_i_* (**A**, **C**). In contrast to the second raw moment, the variance of different growth and death rates overlaps. Differences in death rates mirror the effect of growth rate differences qualitatively. Immigration is *m/N*= 10^−1^, with community size *N* = 10^3^.

## 3 Discussion

Understanding the drivers of communities is one of the main objectives of ecological research. In this work, we have used a stochastic death-birth model with immigration to investigate the equilibrium distribution of communities. Comparing cases where changes only depend on the abundances to cases where types have different birth or death rates, we have identified conditions leading to a robust occurrence-abundance pattern – often reported empirically.

Our approach acknowledges the intrinsic density dependence of communities, Eq. (3), but simultaneously allow us to compute the equilibrium distribution of large and diverse communities, Eq. (7a-7b). Combining a method of model reduction [26] and a detailed balance analysis [24], we asked questions directly linked to empirical observations. In contrast to studies emphasizing biotic interactions [20, 21, 18, 13], our model can be classified with studies that focus on the differential adaptation to the environment [17, 19]. As some of these studies, our results highlight the central role of immigration and biodiversity in community ecology [19, 22].

We tested the reliability of our approach by reproducing known results of neutral adaptation [15]. Namely, that the mean frequency of a type equals its immigration and that the occurrence frequency increases in an S-shaped manner with the mean frequency, Fig. 1. These results already capture the important role of immigration but discard the frequency dependent effects of other types – for which biodiversity might be important.

The match between community level patterns of neutral models and empirical data has been documented extensively [6, 8, 9, 10]. Still, some empirical evidence is at odds with neutral theory [29, 30]. The mismatch with evolutionary history – including phylogenetic trees [29, 30], is one of them. It has been observed that mild differences in adaptation lead to full agreement [31] – indicating the need to consider models with differential adaptation, even if this is mild.

Here, we considered a general death-birth model where large immigration consistently led to a robust occurrence-abundance pattern. Interestingly, evidence suggests that large immigration might indeed be common in various environmental and host-associated microbiomes [10]. Others that deviate from the occurrence-abundance pattern have small immigration [10]. Such seems to be the case in *Caenorhabditis elegans*, where active destruction of microbes during feeding results in reduced immigration to the gut microbiome [32].

A second observation is that with differential adaptation, biodiversity takes a central role. In contrast to the simplest community of two interacting types (Fig. 2-3), diverse communities promote an occurrence-abundance pattern that resembles the neutral case (Fig. 5). With biodiversity, less extreme occurrences and mean frequencies are observed (compare Fig. 2-3 to Fig. 5). Our results agree with research showing that in the limit of high biodiversity, various neutral and non-neutral patterns converge at the community level [19].

Previous research has speculated about the ecological role of types based on their location in the occurrence-abundance curve [10] – the motivation being the possibility to identify microbial taxa actively involved in biotic interactions. Our results indicate that such direct identification from occurrence-abundance curves remains challenging, mainly because neutral and non-neutral types can overlap (Fig. 6). We propose a way forward, based on the inclusion of new observables computed from data [27] (Fig. 7) combined with robust fitting approaches [28].

Our focus at the level of types revealed the difficulty of assessing niche structure and neutrality from empirical data. While niche and neutral patterns can be indistinguishable at the community level, at the level of types, big differences are observed (Fig. 5-6). Commonly, in microbial ecology, models have been tested at the community level, where, embraced by a principle of parsimony, neutral interpretations have been suggested [8, 9, 10]. Our model suggests this is indeed sensible for community level questions. However, for questions at the level of types – including that of ecological roles – general models including differential adaptation can not be avoided. In this case, no parsimonious preference can be given to neutral hypotheses.

The last observation calls for a broader discussion on terminology. As defined by Fisher and Mehta, a community is “statistically neutral” if its distribution can not be distinguished from a distribution constructed under the assumption of ecological neutrality. We must note, however, that ecological neutrality implies statistical neutrality, but statistical neutrality does not necessarily imply ecological neutrality [22]. As our results indicate, a reference to large immigration and biodiversity, rather than neutrality, is more accurate and prevents misleading interpretations, that in their worst form, could lead to unfounded generalizations or hold research questions back. On the contrary, our results suggest that numerous questions about neutrality, adaptation, and ecological roles, in microbial ecology and elsewhere are yet to be answered.

Although we mainly focused on microbial communities, our work can be framed in the larger macro-ecological literature. There, a substantial number of models have linked neutral and niche theories [20, 21, 18, 13, 17, 19]. Heated debates have occurred; however, they have benefited from a close revision of the assumptions on the models and a careful discussion of their implications [19, 6, 31]. The observation of asymptotically equivalent patterns for neutral and non-neutral rates is one of their main results [19]. We believe microbial research can be guided along this line while offering powerful methods to investigate general ecological questions [27]. In particular, the possibility to work, *in vivo* and *in vitro*, with large and diverse communities in much shorter time scales [33].

Finally, we should mention some limitations of our work. A limitation of origin is that we considered a differential adaptation to the environment as the sole source of non-neutrality. Certainly, this is not true in nature, where types take part in numerous symbiotic interactions [13]. Therefore, any empirical application of our model should be preceded by evidence of little to no symbiosis. A technical limitation is that we have only approximated the stochastic dynamics [26]. Our results should be more robust in large communities where types have limited variance [26]. Interestingly, large immigration – which appears to be common in microbial communities [10] – might lead to satisfying this condition.

Although we provided a focused analysis of the occurrence-abundance pattern at equilibrium, future work could study its dynamics [34] and derive exact equations for these and other observables [27]. In addition, identifying neutral and non-neutral types remains an open problem. The development of methods for parameter inference from data [27] seems the way forward.

## 4 Conclusion

Here, we presented a general death-birth model with immigration. Using a method of reduction for the stochastic model, we analysed the equilibrium distribution of abundances for communities equally or differently adapted to the environment. We observe that the community pattern of occurrence-abundance, often reported empirically, is consistently observed in conditions of large immigration and high diversity, regardless of the adaptation to the environment. However, at the level of types, differences in adaptation still lead to large changes.

## Availability of code

The data generated and analysed during the current study can be simulated from the Python code available via GitHub at https://github.com/romanzapien/occurrence-abundance.git.

## Acknowledgements

We thank the *Evolutionary Theory Department* in the MPI Plön and the *CRC 1182: Origins and Functions of Metaorganisms* for fruitful discussions. Finally, we thank the Max Planck Society and the International Max Planck Research School for Evolutionary Biology for the funding provided.

## Authors’ contributions

The original model was developed in discussions between RZC and AT. RZC analysed the model, programmed the code, and wrote the initial draft. All authors interpreted the results, reviewed the manuscript, and approved the final version.

